# Look-up and look-down neurons in the mouse visual thalamus during freely moving exploration

**DOI:** 10.1101/2022.01.22.477320

**Authors:** Patrycja Orlowska-Feuer, Aghileh S. Ebrahimi, Antonio G. Zippo, Rasmus S. Petersen, Robert J. Lucas, Riccardo Storchi

**Affiliations:** Division of Neuroscience and Experimental Psychology, School of Biological Science, Faculty of Biology, Medicine and Health, University of Manchester, UK; Institute of Neuroscience, Consiglio Nazionale delle Ricerche, Milan, Italy

## Abstract

The traditional view that visuomotor integration is a property of higher brain centres has recently been challenged by the discovery in head-fixed rodents that locomotion increases neuronal activity throughout the early visual system (including the retina). Any appreciation of the importance of this behavioural modulation of visual inputs must encompass a comprehensive understanding of the range of behaviours engaged by this mechanism. This information is unavailable from head-fixed preparations in which head and body postures are fundamentally constrained and dissociated from their natural coupling with visual experience.

We addressed this deficit by recording spiking activity from the primary visual thalamus during freely moving exploration, while simultaneously applying frame-by-frame quantification of postures and movements to robust 3D reconstructions of head and body. We found that postures associated with the animal looking up/down affected activity in >50% neurons. The extent of this effect was comparable to that induced by locomotion. Moreover, the two effects were largely independent and jointly modulated neuronal activity. Thus, while most units were excited by locomotion, some expressed highest firing when the animal was looking up (“look up” neurons) while others when the animal was looking down (“look-down” neurons). These results were observed in the dark, thus representing a genuine behavioural modulation, and were amplified in a lit arena.

These findings define the influence of natural exploratory behaviour on activity in the early visual system and reveal the importance of up/down postures in gating neuronal activity in the primary visual thalamus.

## Introduction

A key role of vision is to guide motor actions [1–4]. In turn, motor actions modify vision through self-motion and changes in head and body postures and appropriate interpretation of incoming visual information depends on these parameters [5–7]. For example, mice respond with freezing to a sweeping object flying overhead [2] but with pursuit hunting to a similar object moving at ground level [3], suggesting that selection of the appropriate action requires integration of head and body posture with the visual input. Visuomotor integration has traditionally been studied at high levels of the hierarchical visual pathway (e.g. Posterior Parietal Cortex: [8, 9]; rodent Lateral Posterior Thalamus and primate Pulvinar: [10, 11]). However, it has recently been shown that locomotion (walking/running on a treadmill) is associated with alterations in electrophysiological activity even at the earliest stages of visual processing (including the retina, primary visual thalamus and cortex, and retinal recipient layer of the superior colliculus) [5, 12–17]. There is thus renewed interest in how the incoming visual signal may be modulated by motor actions and the contribution that this makes to visual processing.

A fundamental first step to understanding the mechanisms and functions of visuomotor integration in the early visual system is an appreciation of the range of behaviours contributing to this phenomenon. The head-fixed preparations employed to uncover the influence of locomotion on visual responses [12–20] face a fundamental limitation in this regard as they by necessity constrain movements and dissociate head and body postures from their natural coupling with the visual input. Thus, an important omission in attempts to understand visuomotor integration is the absence of a description of which postures and movements modulate neuronal activity in natural unconstrained conditions. The primary goal of this study was to address this deficit measuring simultaneously postures, movements and firing activity in the primary visual thalamus in freely moving mice.

Success of this endeavour is dependent upon a method to accurately quantify the wide repertoire of postures and movements available to freely moving mice. Computational methods to track body parts in 3D and use these to reconstruct pose at frame-by-frame resolution are increasingly available [21–24]. We have previously developed such an approach suitable for mice [24] and here extend it to measure a wide variety of 3D movements and postures. We find that the higher dimensional description of behaviour enabled by this approach is key to understanding motor influences on the early visual system. Thus, while we confirm that high levels of motor activity (including but not limited to the locomotion available to head fixed animals) excites the primary visual thalamus, we find that head and body postures are at least as influential in defining neuronal activity in this region. Moreover, while movement provides a general increase in firing, the impact of postures is more specific, with a subset of neurons excited by poses characterised by looking upwards (“look-up” neurons), while a different set are excited by poses with associated with looking down (“look-down” neurons). Our discovery, that electrophysiological activity in the primary visual thalamus is influenced by posture during natural exploration, implies that thalamic processing of visual information can be flexibly modulated according to specific visuomotor behaviours.

## Results

### Spontaneous exploration in freely moving animals is defined by independent sets of postures and movements

To capture motor actions in freely moving mice we performed 3D reconstruction of posture and movements in animals implanted with multichannel microelectrodes (see Methods) in dorsal Lateral Geniculate Nucleus (dLGN). During those recordings, animals were spontaneously exploring an open field arena either in the dark or under steady illumination (respectively n = 11, 8 animals). Freely moving animals were recorded in an open field arena by using 4 synchronised cameras (Supp.Fig.1A). Eleven landmarks were identified on the mouse’s body and microelectrode head-stage and were used for tracking the animals (Fig.1A). The tracking data were then triangulated to generate an initial 3D reconstruction of the animal (see Methods). Outliers and missing data in the 3D reconstruction were then corrected by extending a previously developed approach ([24], Methods). The corrected 3D data provided a robust estimate of the animal body over time (Supp.Movie1).

**Figure 1:**
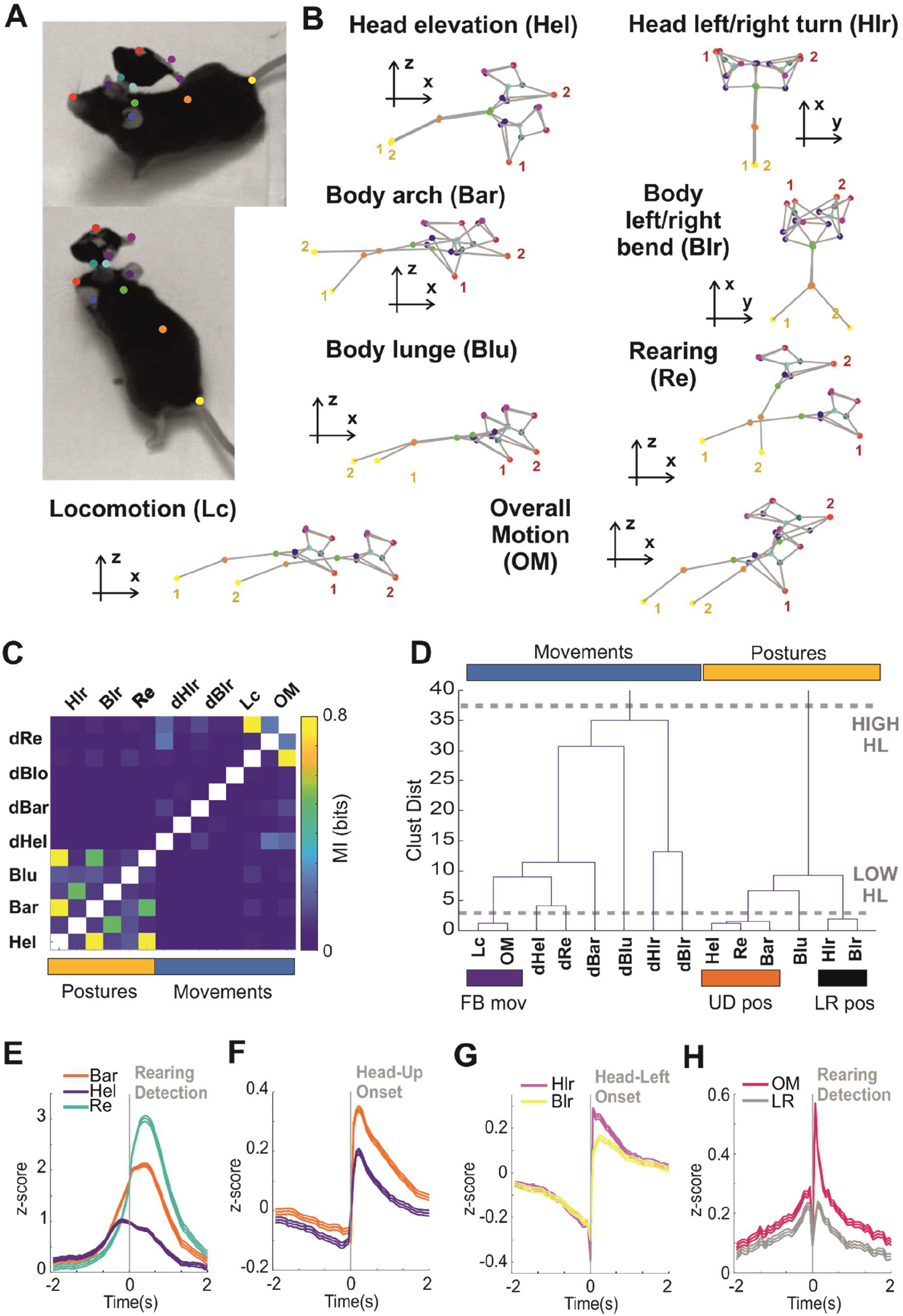
**A)** Visualization of the eleven body landmarks applied to the animal body and the recording head-stage. **B)** Representation of a selection of 8 behavioural state variables. In each vignette we superimpose two poses in which we selectively manipulated one of those variables (to better distinguish the two poses we labelled nose and tail of each pose as “1” and “2”). The axes indicate the viewpoint (x-z for side views and x-y for top views). The other behavioural state variables not represented here (dHel, dHlr, dBar, dBlr, dBlu, dRe) are obtained as temporal derivatives of the associated postures. **C)** Matrix of pairwise MI between behavioural state variables. Postures and movements are indicated by cyan and orange bars. **D)** Hierarchical clustering for all behavioural state variables. Two level of granularities are highlighted (dashed grey lines) indicating respectively a high hierarchy (postures & movements, marked by cyan and yellow bars on top of the graph) and a low hierarchy (full body movements, up/down postures, left/right postures, marked by respectively by purple, orange and black bars). **E)** Average Bar, Re, Hel at the onset of a rearing event (detected at time 0). Note that Bar and Hel precede Re. **F)** Average Bar and Hel at the onset of an upward head movement (detected at time = 0). **G)** Same as panel F but for Hlr and Blr measured at onset of a leftward head movement. **H)** Average OM and Lc at the onset of a rearing event.

From the 3D reconstruction, we quantified the postures and movements of the mouse in terms of a number of behavioural variables (see Supp.Movie2). Hereafter we will define this set of variables as the behavioural state of the animal. Head postures were quantified by head elevation and left/right angles (Fig.1B; Heal elevation: Hel; Head left/right: Hlr). Full-body postures were quantified by projection on the first 3 eigenposes (Fig.1B; Bar: body arch; Blr: body left/right bend; Blu: body lunge), sufficient to explain 80% of changes in the body shape, and by rearing (Fig.1B; Re: Rearing). Six movements were quantified by the temporal derivative of the above postures (Fig.1B; dHel, dHlr, dBar, dBlr, dBlu, dRe). Locomotion was quantified by speed of movement on the x-y plane of the arena (Fig.1B; Lc: locomotion). Finally, we quantified overall motion (Fig.1B; OM: overall motion) by measuring the 3D Euclidean distance between all body points between consecutive frames.

Given biomechanical constraints we expect some postures and movements to correlate with each other. Therefore, we set out to identify relevant groupings among behavioural state variables. To do that we first applied Mutual Information to quantify linear and nonlinear correlations between all pairs. This analysis revealed two main groups that corresponded to postures and movements respectively (Fig.1C). The existence of these two groups was confirmed by a hierarchical clustering analysis (Fig.1D, High HL partition). Hierarchical clustering also revealed that, at a finer grain of analysis, there were three prominent sub-groups (Fig.1D, Low HL partition). (1) Head elevation, body arch and rearing were all associated with animal looking up or down and therefore we defined this group as up/down postures (Fig.1D, UD). (2) Left/right head turn and left/right body bend, that we defined altogether as left/right postures (Fig.1D, LR). (3) Overall motion and locomotion defined hereafter defined as full body movements (Fig.1D, FB). The same sub-groups were observed both in the dark and under ambient illumination (Supp.Fig.1B-E), indicating these sub-groups represent a stable and robust feature of mouse behaviour. Other subgroups were also identified (e.g. dHlr and dBlr, Fig.1D), however these were not consistent across dark and ambient illumination (Supp.Fig.1B-E).

Visual inspection indicated that, although categorization of sub-groups (UD, FB and LR, Fig.1D) was justified on the basis of statistical correlation and similarities in overall behavioural outcome, variables within these sub-groups did not always co-occur (Supp.Fig.2A). Thus, among up/down postures, body arch and head elevation best captured the action of looking up (Supp.Fig.2A-B, respectively poses 2 and 3) and typically preceded full body rearing (Fig.1E). Moreover, changes in head posture could also occur without changes in full body posture. Thus, body arch and body left/right were less strongly coupled than head elevation and head left/right to the onset of vertical and horizontal head movements (Fig.1F-G; Supp.Fig.2A-B, compare poses 5 and 6 for left/right postures). Overall motion was typically dominated by locomotion (Supp.Fig.2A-B, pose 4), but also encompassed vertical actions like rearing (Fig.1H).

### Upward/downward facing postures and full body movements dominate firing rate modulation in visual thalamus in the dark

Having characterised the behavioural state variables involved in spontaneous exploratory behaviour, we then asked whether they covaried with neuronal activity. In order to eliminate the possibility that such correlations could arise from associations between behaviour and visual experience, we first ran these recordings in the dark. A cross-correlation comparison between individual variables and single unit firing rate revealed that a large fraction of units exhibited significant cross-correlation peaks (Fig.3A; shuffle test, see Methods). The most common correlations were with variables encompassing up/down postures and full body movements (Fig.2A, UD & FB). For most variables, a comparable number of units exhibited either positive or negative correlation peaks, while, for full body movements, units exhibited almost exclusively positive correlations (Fig.2A). Across all variables the correlation peaks occurred on average around time zero (see Fig2B), indicating that this aspect of neuronal activity neither systematically predicted nor lagged actions, but rather provided a near instantaneous reflection of the behavioural state. Overall, >70% units (n=69/96) were linearly correlated with up/down and/or full body variables (Fig.2C) and the remaining units had no significant correlations with any other variable.

**Figure 2:**
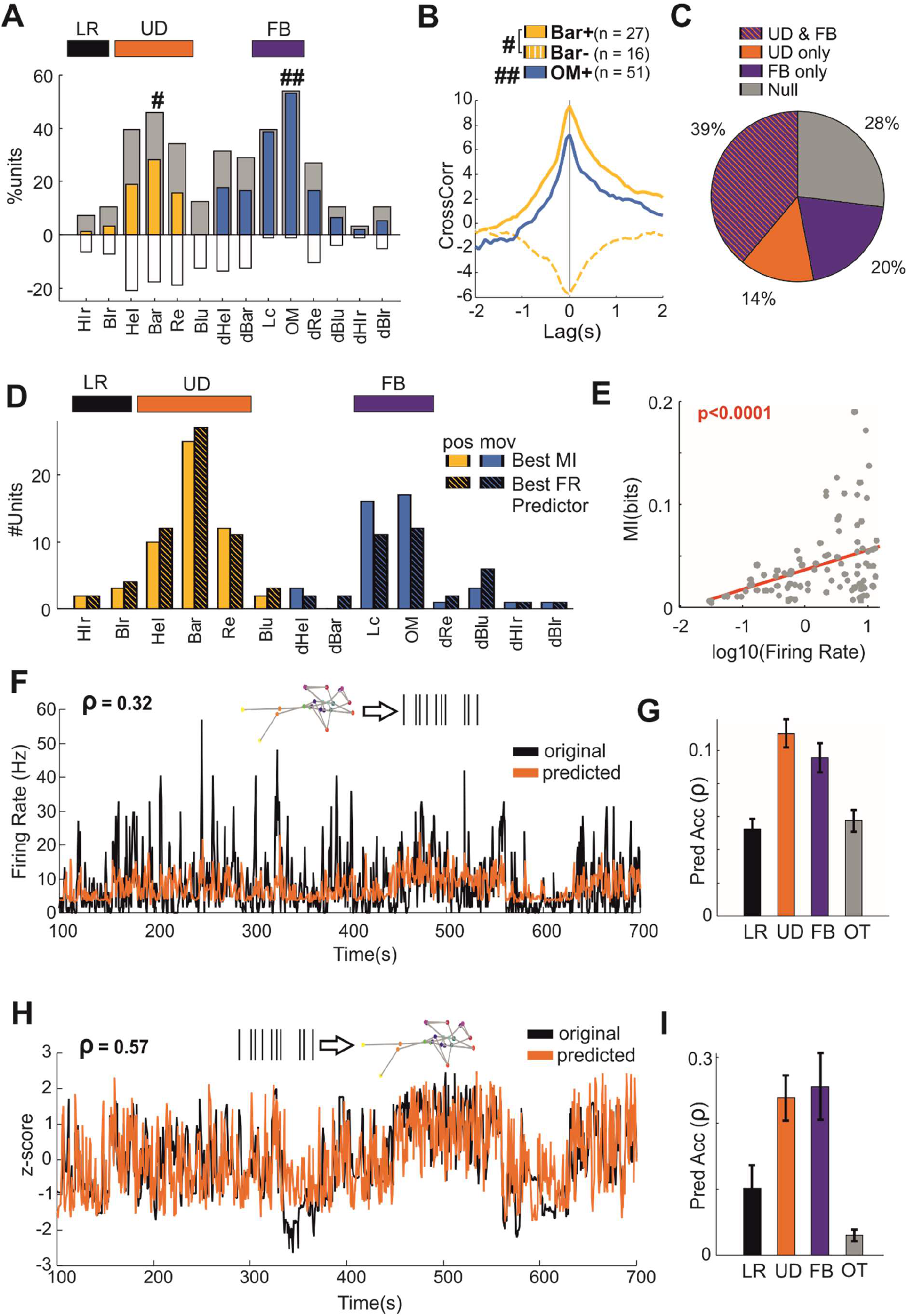
**A)** Results of the cross-correlation analyses. Coloured bar and white indicate the percentage of units with significant positive and negative association with each behavioural state variable. Grey bars indicate the overall percentage of units with a significant association. Full Body movements, up/down and left/right postures are highlighted by coloured bars at the top. **B)** Average cross-correlation across all significant units for Bar (Bar+ and Bar-respectively indicate units with positive and negative peak correlations) and OM. **C)** Percentage of units with significant cross-correlation peak for both UD and FB variables (striped bars), UD only (orange), FB only (purple). The remaining units (OT) have no significant peaks for any of the remaining variables. **D)** Number of units best associated with each behavioural state variable (FB, UD and LR are highlighted by colour bars at the top). Solid colour bars indicate results from MI analyses while striped bars indicate results from prediction analyses. **E)** Mutual Information as function of firing rate (in log units) across all cells (n=96). **F)** Firing rate prediction for a representative unit based on the Bar variable (black = original firing rate; orange = predicted rate). Prediction accuracy, pleasured as Pearson’ correlation between original and predicted is displayed on the top. **G)** Average prediction accuracy (Mean±SEM) for all units as function of LR, UD, FB and OT variables (OT indicates all the remaining 7 variables). **H)** Prediction of Bar based on population firing rates for a representative animal (prediction based on n = 22 units; black = measured Bar; orange = predicted Bar). Prediction accuracy, pleasured as Pearson’ correlation between original and predicted is displayed on the top. **I)** Average prediction accuracy (Mean±SEM) for all animals (n=11) as function of LR, UD, FB and OT variables (OT indicates all the remaining 7 variables).

**Figure 3:**
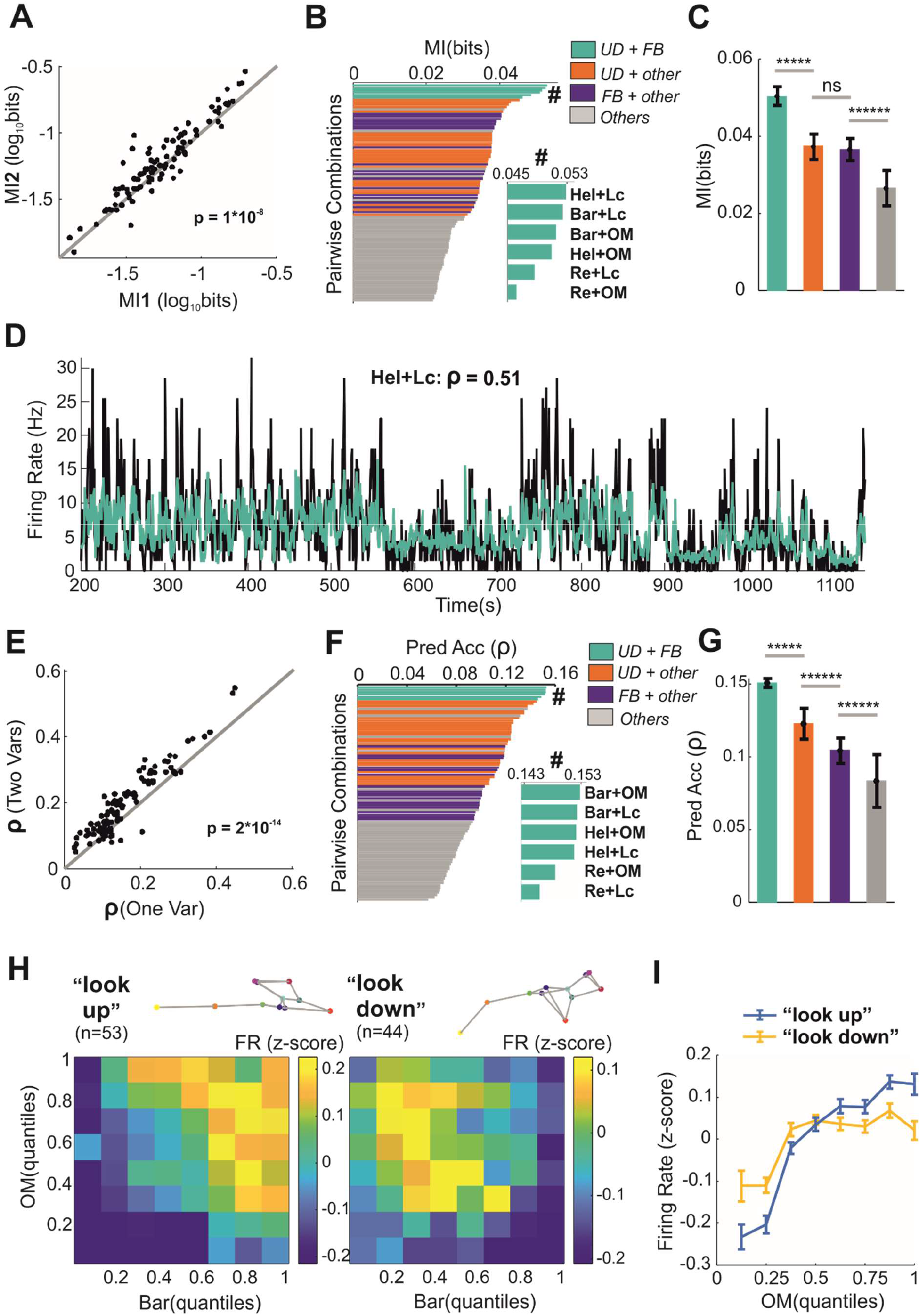
**A)** Comparison between MI conveyed by one predictor (x-axis; MI calculated as shuffle control) and two predictors (y-axis). **B)** Each horizontal bars indicates the average MI between firing rates (n = 96 units) and a pairwise combination of behavioural state variables. Bars are colour-coded according to the type of variables (e.g. green for UD+FB, see legend). The inset (indicated by #) magnifies the top six pairwise combinations. **C**) Mutual Information (Mean±SEM) for each class of predictors. **D)** Firing rate prediction for a representative unit (black = original firing rate; yellow = predicted rate). Prediction accuracy, measured as Pearson’ correlation between original and predicted is displayed on the top. **E-G)** Same as panels **(A-C)** but here we used prediction accuracy instead of Mutual Information. **K**) Z-score transformed firing rate (color coded) as function of body arch (Bar) and overall motion (OM). The units are divided into “look-up” and “look-down” units. **I**) Z-score transformed firing rates (mean±SEM, n = 52, 45, respectively look-up and look-down units) as function of overall motions (OM).

In order to capture both linear and nonlinear correlations between motor state variables and single unit activities we applied Mutual Information (MI). For each unit we then identified the single variable with highest MI (corresponding to that unit’s ‘favoured’ behavioural variable). Most units (83% cells, n = 80/96) favoured variable was either with an up/down posture or a full body movement (Fig.2D, solid colour bars). Among up/down postures, units conveyed higher information for body arch than for rearing (p = 0.0056, sign = 62, n = 96, sign-test), while body arch and head elevation were not significantly different (p = 0.1253, sign = 40, n = 96, sign-test). Among full body movements, information conveyed by locomotion and overall motion were also comparable (p = 0.36, sign = 53, n = 96, sign-test). Across the recorded neuronal population, the amount of information was proportional to the firing rate, so that units with higher firing rates carried higher information (Fig.2E, p<0.0001, n = 96, t-test). These results indicate that the modulation of dLGN neurons by posture mainly reflected sensitivity to body arch, which encompasses head elevation (Fig.1B; Supp.Movie 2), while modulation by movements was dominated by locomotion.

Since single units convey information about behavioural state variables, we would expect that such variables would predict single neuron activity to a significant extent. To address this possibility, we trained a model (based on XGBoost, [25], see Methods) to predict firing rate from individual behavioural variables. We then evaluated their prediction accuracy by calculating Pearson’s linear correlation between recorded and predicted spike-counts on crossvalidated data. Based on these results, for each unit we identified the best predictor, i.e. the variable associated with highest Pearson’s correlation. Across our dataset the best predictors captured substantial amounts of variability (ρ = 0.16±0.08, mean±SD; see representative unit in Fig.2F and Supp.Fig.3A). To exclude the possibility that this result was an artifact of our analyses, we used the same variables to generate shifted predictors that preserved structure of the original predictors but were no longer associated with neuronal activity (see Methods). The accuracy of shifted predictors was consistently disrupted across our dataset of recorded units (Supp.Fig.3B). Overall, prediction accuracy for the original behavioural state variables returned results that were consistent with those obtained by using Mutual Information. Thus, up/down postures and full body movements were the best predictors for 76% of the units (Fig.2D, striped bars), body arch was the best predictor among postures and significantly surpassed rearing (p = 0.0056, sign = 63, n = 96, sign-test), while, among movements, the best predictions provided by overall movement were not significantly higher than locomotion (p = 0.2615, sign = 54, n = 96, sign-test). Among sub-groups, both up/down postures and full body movements provided better predictions than left/right postures and all other variables (Fig.2G; p<0.0005 across all comparisons, sign-test, n = 96). Similarly to MI results, prediction accuracy was also proportional to firing rate (p<0.0001, n = 96, t-test).

Finally, having shown that single unit firing can be significantly predicted by behavioural state, we asked whether the opposite was also true: can we predict be behavioural state variables from population activity? To test this, we trained an algorithm (random forests [26], see Methods) to predict individual behavioural state variables based on spike counts from all units that were simultaneously recorded from each animal. Consistent with previous results, we found that up/down postures and full body movements were the best predicted variables (Fig.2H-I; p = 0.0002, χ^2^ = 19.22, n = 11 animals, Kruskal-Wallis test), providing respectively 135% and 152% increase in accuracy over left right postures and 692% and 749% over other variables.

Overall, these results show that, during spontaneous exploration of mice in the dark, neurons in visual thalamus are modulated by behavioural state and the effect is largely associated with upward/downward facing postures and full body movements.

### Single unit firing is jointly modulated by upward/downward facing postures and full body movements in the dark

Does each individual unit encode a single behavioural variable or does it encode more than one? To answer this we first estimated Mutual Information between single unit activity and all possible pairs of variables. For each unit we then selected the pair associated with the highest information. We first asked whether pairs of variables provided more information than individual ones. To provide the fairest comparison we re-estimated MI for individual variables by using pairs in which the values from one variable were shuffled over time in order to remove its association with neuronal activity (see Methods). We found that most units conveyed more information about variable pairs than about individual variables (Fig.4A, p = 1*10^-8^, sign-test, n = 96). We next asked which variables provided the most effective modulations. We found that pairwise combinations of up/down postures and full body movements were the strongest modulators (Fig.3B), providing, on average, a 90% increase over other pairwise combinations that did not include those variables (Fig.3C).

**Figure 4:**
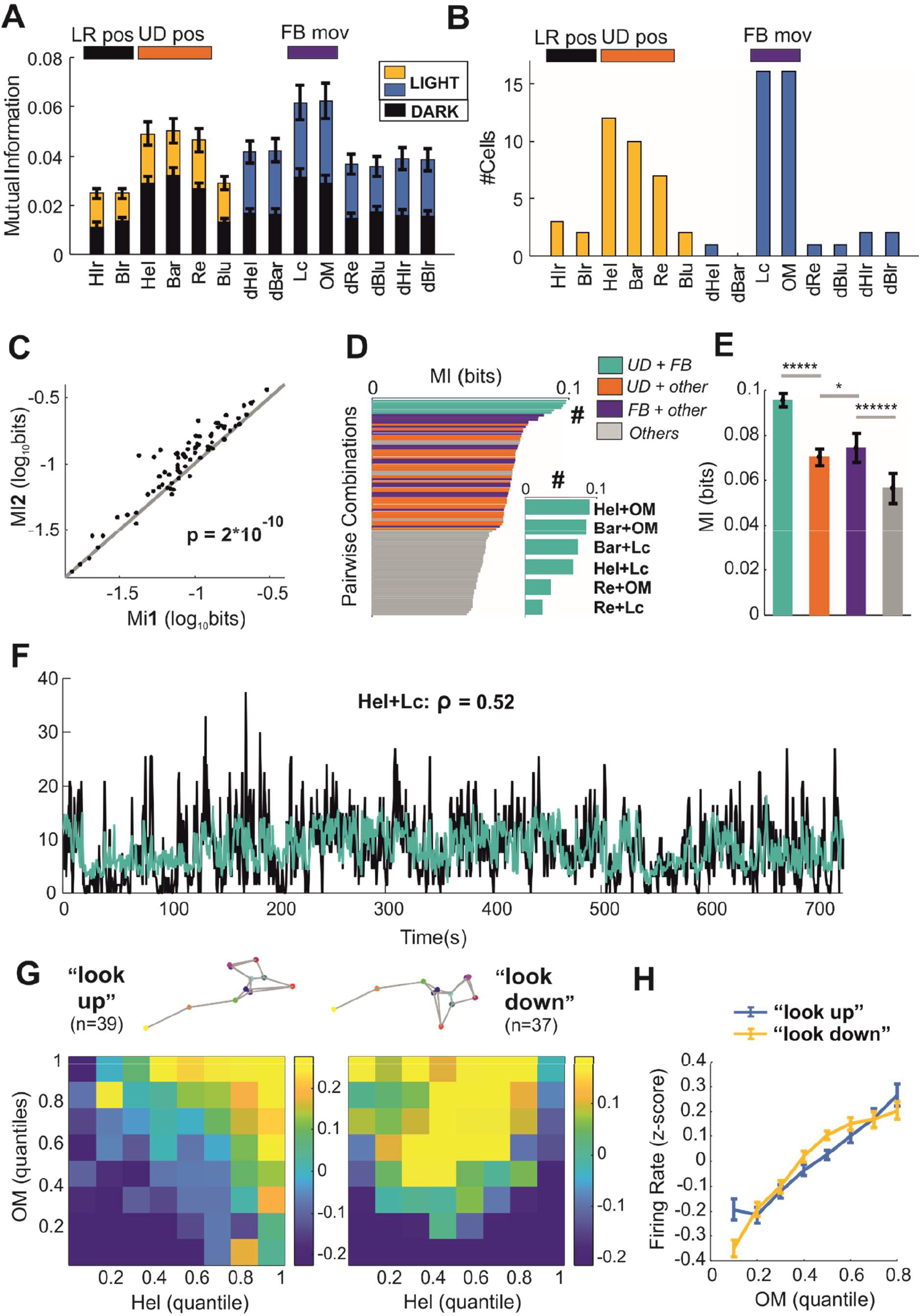
**A)** Mutual Information (mean±SEM) for each motor state variable in dark and illuminated environments. **B)** Same as **Figure 2D** but for animals recorded in an illuminated environment. **C-F)** Same as **Figure 3A-D** but for animals recorded in an illuminated environment. **G-H)** Same as **Figure 3H-I** but for animals recorded in an illuminated environment.

We then asked whether pairs of behavioural state variables, when compared with individual variables, could provide better predictions of neuronal activity. We found that this was the case, since for most units Pearson’s correlation between predicted and original activity increased when using two predictors (Fig.3D-E; p = 2*10^-14^, sign-test, n = 96). Like for MI analyses, pairwise combinations of up/down postures and full body movements were the best predictors (Fig.4F), providing an average 81% increase over pairwise combinations that did not include those variables (Fig.4G).

Finally, we asked which and how many different types of modulations could be observed at single unit level. To answer this question we performed an unsupervised clustering analysis (based on community detection [27], see Methods). Each unit was represented as a 2D histogram of the average firing rates as function of body arch and overall motion (the two strongest modulators). The clustering process then automatically determined the number of distinct types. This analysis revealed two classes of units: “look-up” units, most active when the animal assumed an upward facing posture (Fig.3H, left panel, n = 53) and “look-down” units, most active when the animal assumed a downward facing posture (Fig.3H, right panel, n = 44). Both “look-up” and “look-down” units were positively modulated by increased levels of motor activity (Fig.4I; look-up: p = 1*10^-37^, 8*10^-12^, χ^2^ = 190.12, 66.4; n = 52, 45; Kruskal-Wallis test for look-up and look-down units). Similar clustering results were obtained by combining overall motion with head elevation or rearing (Supp.Fig.3C-D).

Overall, these results show that modulation of single dLGN units is a multi-dimensional effect in which two distinct populations, both excited by high motor activity, are recruited according to the mouse up/down postures.

### Modulation of single unit activity by behavioural state is maintained and amplified by ambient illumination

All modulations described so far were obtained by recording the animal in the dark, i.e. in absence of visual stimulation. We next asked whether these modulations persist when the visual thalamus is additionally stimulated by visual inputs. To do that we repeated our freely moving experiments but this time with the arena illuminated (within the photopic range).

In the light arena average values of MI across all postures and movements were higher than in the dark (Fig.4A, p<0.005 for all variables, rank-sum tests, n = 96, 75 for dark and light conditions). This was not simply a reflection of higher firing as firing rate was not significantly different between light and dark conditions (p = 0.86, rank-sum = 8200.5, zval = −0.17, n = 96(dark), 75(light), rank-sum test). All other results were qualitatively recapitulated. Up/down postures and full body movements were still associated with largest MI values in the majority of the cells (81%, n = 61/75, Fig.4B). Consistently with recordings in the dark, most units conveyed more information about pairs of behavioural state variables than about individual variables (Fig.4C, p = 5*10^-11^, sign = 65, n = 75). Again, pairwise combinations of up/down postures and full body movements were the strongest modulators (Fig.4D), providing, on average, a 70% increase over other pairwise combinations that did not include those variables (Fig.4E). Results based on predictive modelling approach were consistent with MI analysis (see representative unit in Fig.4F and population results in Supp.Fig.4A-E). Clustering analysis revealed two classes of units that qualitatively matched the “look-up” and “look-down” units observed in the dark (Fig.4G) and both classes were similarly modulated by full body movements (Fig.4H, p = 2*10^−22^, 4*10^-31^; χ^2^ = 117.78, 159.75; n = 39, 37; Kruskal-Wallis test for look-up and look-down units). Finally, population activity predicted up/down postures and full body movements with higher accuracy compared with other motor state variables (Supp.Fig.4F-G).

These results show that the main properties of the modulations described in the dark are maintained and amplified with visual stimulation.

## Discussion

In everyday life, visual processing is concurrent with motor actions. It follows that to understand visual processing, we also need to understand how specific actions affect neuronal activity along the visual pathway. Previous studies found strong modulations of neuronal activity by motor activity along the visual pathway and throughout the brain [14, 16–19, 28–30]. However, since most studies were performed in head-fixed animals, it is still unclear which aspects of behaviour would provide the relevant modulations in natural freely moving conditions. Our study aimed to fill this gap in the mouse dLGN, the key subcortical station linking the retina to the primary visual cortex.

Our first result was that upward and downward facing postures affected firing rate of >50% of neurons in dLGN. Encoding of head and body postures was previously described in the rat posterior parietal cortex and frontal motor cortex [31] but, to the best of our knowledge, not in the visual system. Most studies on visual processing focussed on the effect of locomotion on a treadmill and revealed that this behaviour substantially affect neuronal activity in primary visual cortex (see e.g. [17–19, 30, 32]), and dLGN and LP regions of the visual thalamus [14, 16, 20]. Since those studies were performed in head-fixed animals, the nature this preparation did not allow for measuring head movements and head and body postures. Two recent studies, performed in freely moving animals, quantified the effect of head movements but not of head and body postures [4, 5]. Our results indicate that the effect of head movements (e.g. left-right head turns) on firing rates is present in dlGN but significantly less prominent than the effect of upward and downward facing postures (see e.g. Fig.2G,I).

Our second result was that modulation exerted by upward and downward facing postures was largely independent from, and interacted with, the modulation exerted by full body movements (and typically locomotion). This result addresses a long-standing debate about the nature of behavioural modulation in dLGN. Neuromodulation of primary visual thalamocortical loops has been traditionally associated with the control of sleeping and arousal states [33]. More recent studies have shown that modulation of neuronal activity in these regions is related to both motor activity and arousal (as measured via pupil dilation) [12–14, 17, 34, 35]. Separating the arousal component from motor activity component has been traditionally difficult using head-fixed preparations since locomotion on a treadmill is strongly coupled to pupil dilation [13, 14, 35, 36]. Thus, while intermediate levels of arousal can occur without locomotion [12, 32, 37, 38], locomotion always coincide with high pupil dilation [20, 34, 38]. Our experiments in freely moving animals reveal two largely independent modulations, respectively by full body movements and upward and downward facing postures. Thus, while locomotion evoked a generalised increased in firing rate likely associated with high arousal, this effect was gated by upward and downward facing postures, so that some units were most active when animals pointed their head high (“look-up” units), while others when animals kept their head low (“look-down” units). This result indicates that neuronal modulation in dLGN can be behavioural-specific (e.g. some neurons will be active while exploring the ground, others while searching the sky) rather than simply reflecting a generalised state of arousal or motor activity. Additional experiments in other primary sensory thalamic regions will be required to understand whether these results also apply to other sensory modalities.

Our final result was that, when experiments were repeated in an illuminated environment, modulations by postures and movements were maintained and amplified. The most parsimonious explanation is that the amplification would arise from the introduction of movement-related visual stimuli produced by self-motion. Alternatively, ambient light could drive neuronal activity of the visual thalamus into a more excitable state [39–42] and this could amplify modulations observed in the dark. Finally, if behavioural modulation is inherited from the cortex, ambient light could modify the interactions between excitatory and inhibitory populations in primary visual cortex [43, 44] and, in turn, the corticofugal feedback onto visual thalamus.

The sources of modulations by the behavioural state on the primary visual thalamus are currently unknown. Modulation by upward and downward facing postures could be provided by vestibular afferents from the brainstem, since optogenetic stimulation of the Medial Vestibular Nucleus diffusely excites sensory thalamic nuclei and cortices [45]. Consistently with this possibility, results from anaesthetized cats showed that visual responses of >%80 neurons in dLGN were modulated by electrical stimulation of vestibular nuclei in the brainstem [46]. Primary visual cortex could also play an important role, by conveying both postural and motor information via the extensive direct and indirect (via Thalamic Reticular Nucleus, TRN) corticofugal projections [47]. Strong inputs from thalamic regions other than TRN are unlikely given the sparse intra-thalamic connectivity [48]. Visual thalamus has also been shown to receive afferents from the superior colliculus [49] and parabigeminal nucleus [50], two important regions involved in visuomotor behaviours and action selection [51]. Additionally, direct or indirect input from the mesencephalic locomotor region (Pedunculo Pontine Nucleus, Laterodorsal Tegmental Nucleus) could be involved [50, 52]. Finally, recent studies also provided evidence that arousal and locomotion modulate input from the retinal afferents [12, 13] and this modulation affects sensory processing in visual thalamus. Further studies will be needed to investigate the relative contribution of those sources.

Our results indicate that most neurons in visual thalamus are modulated by multiple components of the ongoing behaviour. This is consistent with recent studies of brain-wide modulation (for a review see [53]), indicating that spontaneous neuronal activity is high dimensional and captures many distinct components of spontaneous behaviour. We currently do not understand the functional role(s) of such a widespread representation of behaviour. One possibility is that excitation of specific subsets of neurons would provide a flexible encoding scheme to re-purpose visual processing according to ongoing behaviour. Indeed, locomotion has been shown to modulate the gain of visual responses [17, 54] and this modulation can selectively amplify specific visual features (e.g. transient ON responses [20]). Here we show that, in a freely moving animal, modulation is richer than just locomotion, and different neurons are associated diverse visuomotor contingencies, indicating higher levels of flexibility. Consistent with our results, eye movements, largely suppressed in head-fixed preparations, are strongly driven by changes in upward and downward facing postures in freely moving animals [55]. The richer modulations observed in freely moving conditions could also be employed to support coordinate transformation and spatial navigation during spontaneous exploration [19, 56] or to learn new visuomotor contingencies by gating visual inputs according to behavioural context [57]. Finally, behavioural modulation could part of an encoding scheme that predicts incoming visual inputs based on self-motion [18]. Altogether, these hypotheses are not mutually exclusive, and the outputs of the visual system could be employed by the brain in more than one way.

In summary, this study fills a gap in our understanding of how behaviour modulates the early visual system in natural unrestrained conditions. Our results indicate that neuronal activity in primary visual thalamus can be flexibly modulated according to movements and specific postures. The extent to which these modulations are applied to different stages of the visual pathway (and to other sensory pathways) is largely unknown. We believe that further investigation on this topic constitutes an important avenue for future studies.

## Methods

### Animals

Experiments were conducted on 11 adult, male C57BL/6J mice (Charles River). Experiments were conducted in accordance with the Animals, Scientific Procedures Act of 1986 (United Kingdom) and approved by the University of Manchester ethical review committee.

### Recovery Surgery

Throughout the procedure mice were anaesthetised with isoflurane (95/5% Oxygen/CO2 mix; flow rate: 0.5 – 1.0L/min). Concentrations of 4 – 5% and 0.5 – 1.5% were used respectively for induction and maintenance of anaesthesia. The level of anaesthesia was verified by the lack of withdrawal reflex. During the surgery animal’s body temperature was automatically maintained at 37°C by a heating mat and animal’s eyes were protected from drying out by eye drops. Once placed in the stereotaxic frame (Narishige, Japan), the animal fur was trimmed and 1% EMLA cream applied topically to the surrounding skin. An incision was made to expose a skull surface and set stereotactic *bregma* point, craniotomy and screws coordinates. Next, two slotted cheese machine screws (M1.6×2.0mm, Precision Tools, UK) were inserted respectively into the parietal, and interparietal plates to act as anchors for the dental cement and for electrical grounding of the electrode. After craniotomy the electrode was inserted into the dLGN (coordinates from *bregma*: 2.0-2.5mm medial-lateral, 2.3-2.5 mm rostro-caudal) at a depth of 2.8mm from the brain surface. Light-curing cement (X-tra base, VO64434-A, VOCO) was applied to seal the implant. After surgery, the mouse was released from the ear bars and allowed to recover in a single-housed heated cage. Analgesia was provided with an intramuscular injection of 0.05mg/kg buprenorphine. After the procedure the animal was allowed to recover in a single-housed home-cage for a minimum of six days prior to experimentation.

### Experimental Set-Up

A detailed description of the experimental set-up for behavioural recordings is provided in [58]. The animals were recorded in a square open field arena (dimensions: 30cm x 30 cm, Supp.Fig.1A). Behavioural recordings were acquired with 4 programmable cameras (Chamaleon 3 from Point Grey; frame rate = 15Hz) equipped with infrared cut-on filters (Edmund Optics) to isolate light in the infra-red range. Neuronal recordings were performed via 16 channel electrodes (Neuronexus; model: A4×4-3mm-50-125-177; package: CM16) with a TBSI W16 wireless acquisition system (Triangular BioSystems; sampling rate = 30 kHz). Frame acquisition, controlled via Psychopy (version 1.82.01, [59]), was synchronized with acquisition of neuronal data via an Arduino Uno board (www.arduino.cc). This board delivered a common electrical trigger to the cameras and the electrophysiological acquisition system.

### Behavioural Protocol

Naïve animals were briefly anaesthetised (~2 minutes) with 3% isoflurane in order to connect the electrode head-stage. They were gently positioned at the centre of the arena and allowed plenty of time to recover from the anaesthesia. After the animals expressed sustained exploration of the arena (typically ~20 minutes after being placed in the arena) the experiment started. Animals were not specifically trained and freely explored the arena throughout the duration of the experiment (typically ~45 minutes). During recordings in the dark, brief steps of light (0.66-1.33 seconds) were provided to test for visual responses. During recordings under steady illumination (4.08*10^10^, 1.65*10^13^, 1.94*10^13^ and 2.96*10^13^ photon/cm^2^/s respectively S-cone opsin, Melanopsin, Rhodopsin and M-cone opsin) brief steps of dark (0.66-1.33 seconds) were also delivered. Video recordings were performed in epochs of 15-24 seconds, each separated by 30-40 seconds.

### Reconstruction of 3D Poses

An initial 3D reconstruction of the mouse body was obtained by triangulating body landmarks from the four cameras (see body landmarks in Fig.1A). The four-camera system was calibrated as previously described [24]. Tracking of body landmarks from individual cameras was performed with DeepLabCut [60]. The algorithm was trained with ~1000 manually annotated images and ran on a dedicated Ubuntu machine equipped with a Titan RTX GPU (Nvidia, Santa Clara, California, USA). The initial 3D reconstruction was contaminated with missing data and outliers, typically due to occluded body points. In order to correct the 3D reconstruction we modified a previously developed algorithm [24]. We first estimated a Statistical Shape Model (SSM, [61]) from a dataset of 350 manually re-annotated 3D poses. The poses were initially aligned by used Procrustes Superimposition (with scale parameter = 1). The SSM was then estimated as by applying Probabilistic Principal Component Analysis (PPCA, [62]; MATLAB function: *ppca*). The SSM allowed us to express the 3D pose of the animal position as

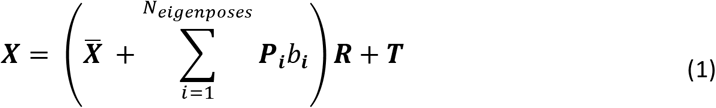

Where: ***X*** is an *N_p_ x*3 matrix representing the 3D pose for given frame (*N_p_* is the number of body landmarks); 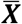 is the mean pose; ***P_i_*** and *b_i_* represent respectively the *i^th^* eigenpose obtained by training the SSM and its score (the shape parameter); ***R*** and ***T*** are respectively the 3 *x* 3 rotation and *N_p_ x*3 translation matrices that map the animal’s position in the experimental environment. Note that is *T* obtained as ***T*** = ***t*** 0 1 where ***t*** = [*t_x_, t_y_, t_z_*] is the 3D translation vector and **1** is an *N_p_ x* 1 all-ones vector. To obtain a robust 3D reconstruction we minimised, as function of (***b, R, T***), the following cost function

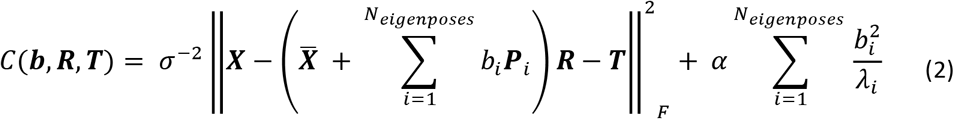

Where: *σ*^2^ represents the noise term obtained from the PPCA, *λ_i_* the eigenvalue associated with the *i^th^* eigenpose and *α*, the regularization parameter, was set at 0.01. Outliers in a pose were flagged when the *C* > *inv*(*X*^2^(3*N_p_*)), with *N_p_* indicting the number of body landmarks. When this happened, we removed the body landmark associated with largest value of *C*, reduced *N_p_* by 1, and recalculated *C*. This operation was performed iteratively until *C* < *inv* (*X*^2^(3*N_p_*)). Then, in order to fill-up missing values and correct the remaining 3D coordinates, we recalculated ***X*** as in equation (1), by using (***b, R, T***) values obtained from the minimization of equation (2) and 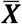, *P* provided by the SSM.

### Quantification of Motor State Variables

Quantification of motor state variables was based on 3D data after applying the reconstruction algorithms described above. Head elevation (Hel) was calculated as the angle between the nose and the neck landmarks. Head left and right turns (Hlr) as the angle between midline and nose. Body arch (Bar), body left and right turns (Blr), and body lounge (Blu) corresponded to the shape parameters associated respectively with first, second and third eigenposes. These definitions (e.g. body arch) were based on visual inspection of the movies in which we visualised the full body changes in shapes along each eigenpose (see Supp.Movie2). Rearing corresponded to the z-coordinate of the translation matrix ***T***. Locomotion (Lc) was calculated as Euclidean distance on the x-y axis of the body centre between two consecutive frames. The body centre in each frame was defined by the values of the translation vector ***t*** (see above). Overall motion (OM) was defined as the Euclidean across of all body points between two consecutive frames. All motor state variables where then transformed into z-scores for all further analyses.

### Hierarchical Clustering

Hierarchical agglomerative clustering across all motor state variables was applied to the Mutual Information matrix (Fig.1D). The hierarchical trees were created by using nearest distance method (MATLAB function: *linkage,* metric: *weighted*).

### Clustering of Firing Rate Distributions

In order to cluster firing rate as function of two motor state variables (see Fig.3K and Fig.4I) we used community detection as described in [27]. The only free parameter of this algorithm (γ) was set to 1. The adjacency matrix was calculated by using pairwise Spearman’s correlation between units. The results of this calculation were binarized by using the median as median of the original matrix as threshold.

### Pre-processing of Neuronal Data

Action potentials (typically >50 μV, see Supp.Fig.5A) were extracted from the continuous, high-pass filtered signals (low cut-off at 250 Hz) by using Offline Sorter (version 3). Noise artifacts were then removed, and individual units were sorted using the template sorting (Valley Seek) method and Principal Component Analysis (PCA) in Offline Sorter software (Plexon, USA) and next manually inspected. Reliable single unit isolation was confirmed by referenced to MANOVA F statistics, J3 and Davies-Bouldin validity metrics (Offline Sorter). Signal-to-noise ratio (SNR) was then calculated for each unit. Only units with SNR>1.5 were kept for further analysis (Supp.Fig.5B-C). In order to avoid double counting the same unit from different channels we merged units that shared > 50% spikes timestamps from pairs of neighbouring channels (this only happened twice).

### Cross-correlation Analyses

Cross-correlation between spike counts and motor state variables was estimated after removal of the mean from each signal. In order to test for statistical significance we estimated a null-distribution by using a shuffling procedure. The association between spikes and motor state variables was broken by dividing spike counts into epochs and randomly permuting the order of such epochs. This operation was repeated 1000 times in order to generate the null-distribution. Original cross-correlation was deemed significant when at least one of its value (range [−2,+2] seconds; time-bin = 0.0667 seconds) was outside a [0.0005,0.9995] confidence interval.

### Mutual Information Analyses

Prior to estimation, continuous motor state variables were transformed into discrete variables by applying quantile discretization. Probability distributions were then estimated from the frequency histograms of each signal. These distributions were used to obtain response and noise entropies, respectively calculated as

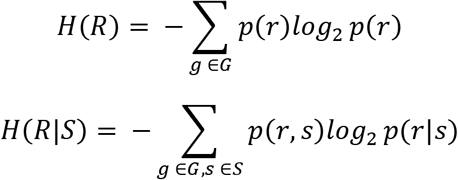

Where: *r* indicates spike counts and s indicates the values of the motor state variables. Both entropy terms were corrected for the sampling bias by using quadratic extrapolation as in [63]. Shannon’s Mutual Information (MI) was then calculated as *MI* = *H*(*R*) – *H*(*R*|*S*). When comparing MI obtained from one vs two motor state variables, we adopted the following strategy. First MI was estimated for two motor state variables as described above. Then, MI for single variables was estimated from the same two variables but by previously shuffling the order of one or of the other variable. Finally, the largest MI obtained by shuffling was taken as estimate of MI for an individual variable.

### Predictive Modelling Analyses

Prediction of spike counts based on motor state variables was performed with Extreme Gradient Boosting (XGBoost, [25, 64]; parameters: learning rate = 0.025; number of boosting iterations = 500; evaluation metric: log-likelihood loss; subsample = 1; maximum depth = 3; gamma = 1; tree method: gpu_hist). This method has been shown to outperform spike count predictions obtained with more standard approaches based on Generalized Linear Models [65]. Prediction of behavioural state variables was based on random forest [26] (parameters: maximum number of splits = 100; number of cycles = 100). For this latter analysis the variables were discretized into 100 bins. All datasets were bisected into two equal parts that were used for training and cross-validation. Prediction performance on the cross-validation set were measured as Pearson’s linear correlation between predicted and original firing rate. For prediction of spike counts, in order to test for the possibility that positive correlations were obtained purely by chance, we repeated this analysis by shifting the behavioural state variables. Shifting was obtained by splitting the behavioural state time series into two equally long epochs and inverting the order of those epochs. In this way the association of these time series with spike counts was abolished while the temporal structure of those series was preserved.

## Acknowledgements

We wish to express our gratitude to James Morizio and Harry Fu for support with TBSI recording system and to Annette Allen for comments and advice on the manuscript. This study was funded by: a Sir Henry Dale fellowship from Wellcome Trust to R.S. (Grant Code: 220163/Z/20/Z); a David Sainsbury Fellowship from National Centre for Replacement, Refinement and Reduction of Animals in Research (NC3Rs) to R.S. (Grant Code: NC/P001505/1); the European Union’s Horizon 2020 research and innovation programme under the Marie Sklodowska-Curie grant to P.O.F (Grant Code: 897951); a Bekker Programme grant implemented by the Polish National Agency for Academic Exchange to P.O.F. (Grant Code: PPN/ BEK/2018/1/00192); a Wellcome Trust Investigator award (210684/Z/18/Z) to R.J.L; a research grant by Biotechnology and Biological Sciences Research Council to R.S.P. and R.S. (BBSRC; Grant Code: BB/V009680/1); a grant from Weizmann – UK Making Connections Programme to R.S.P.

---

**Supplementary Figure 1:**
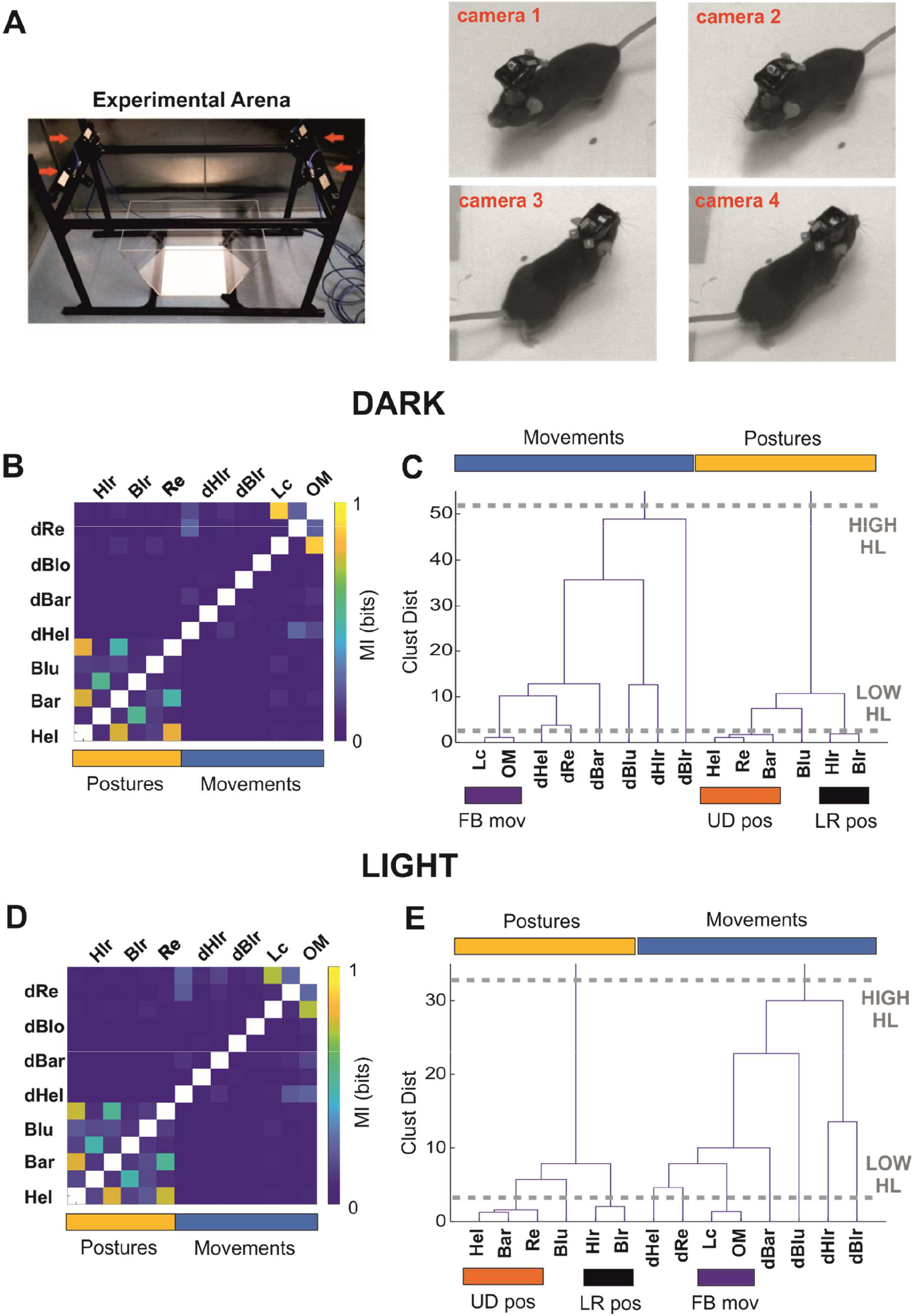
**A)** A pictures of the experimental apparatus (left panel). The arena is placed in the centre and imaged by four overhead cameras (indicated by red arrows). A representative frame simultaneously acquired by the 4 cameras is shown (right panel). **B-C)** Same as Figure **1C-D** but for animals recorded in the dark. **D-E)** Same as Figure 1C-D but for animals recorded under photopic light.

**Supplementary Figure 2:**
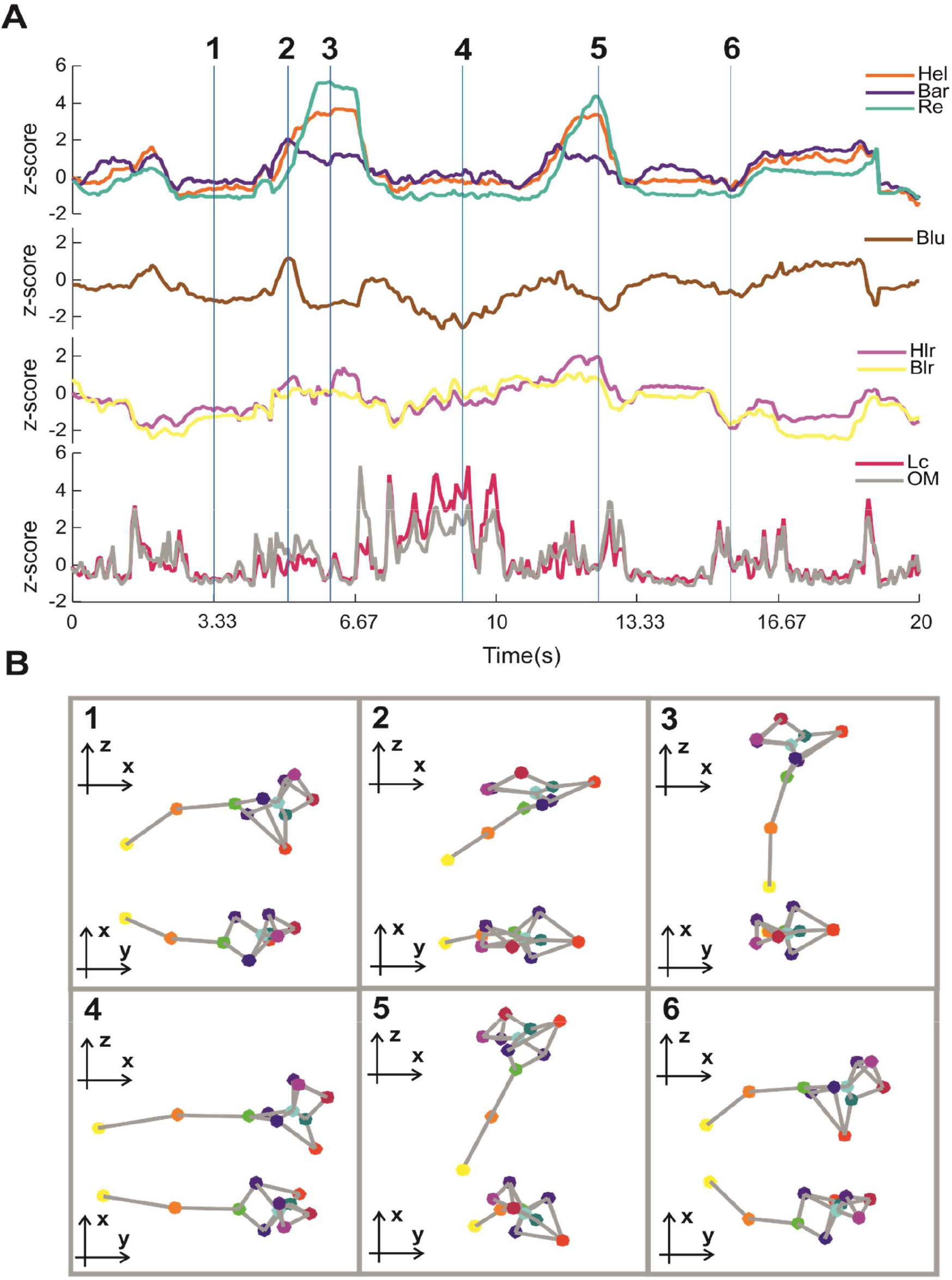
**A)** Representative time series for a selection of behavioural state variables (Hel, Bar, Re, Blu, Hlr, Blr, Lc, OM - the variables not shown are simply obtained as time derivative of the postures). **B)** Poses from six frames (indicated in panel A by vertical blue lines) are shown from side and top view.

**Supplementary Figure 3:**
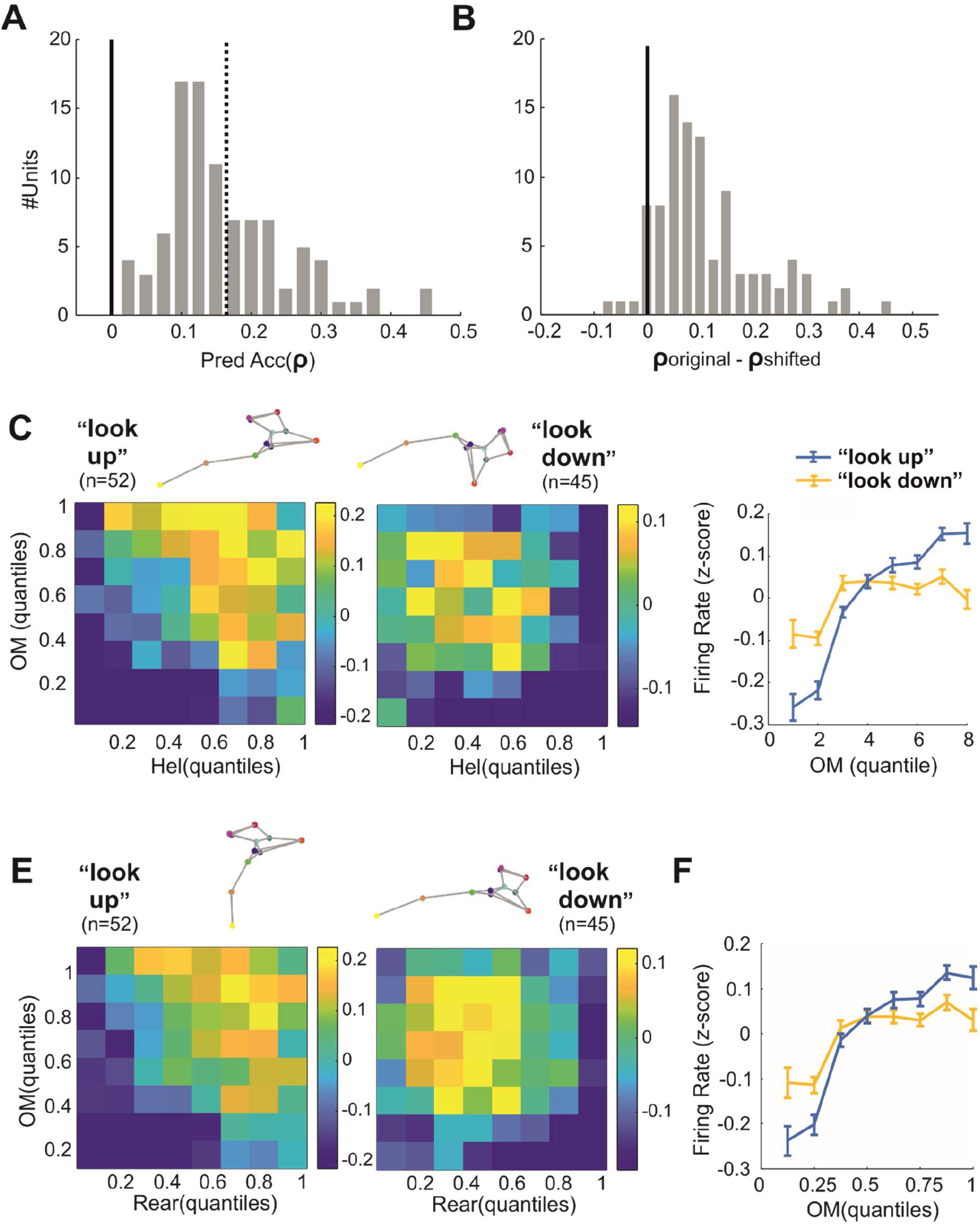
**A)** Distribution of prediction accuracy (measured as Person’s p) for the dataset of units recorded in the dark (n = 96). Dashed line indicates average accuracy. **B**) Paired difference in prediction accuracy between original and shifted predictors (n = 96). **C-D)** Same as Fig.3H-I but for combinations of Hel and OM. **E-F)** Same as Fig.3H-I but for combinations of Re and OM.

**Supplementary Figure 4:**
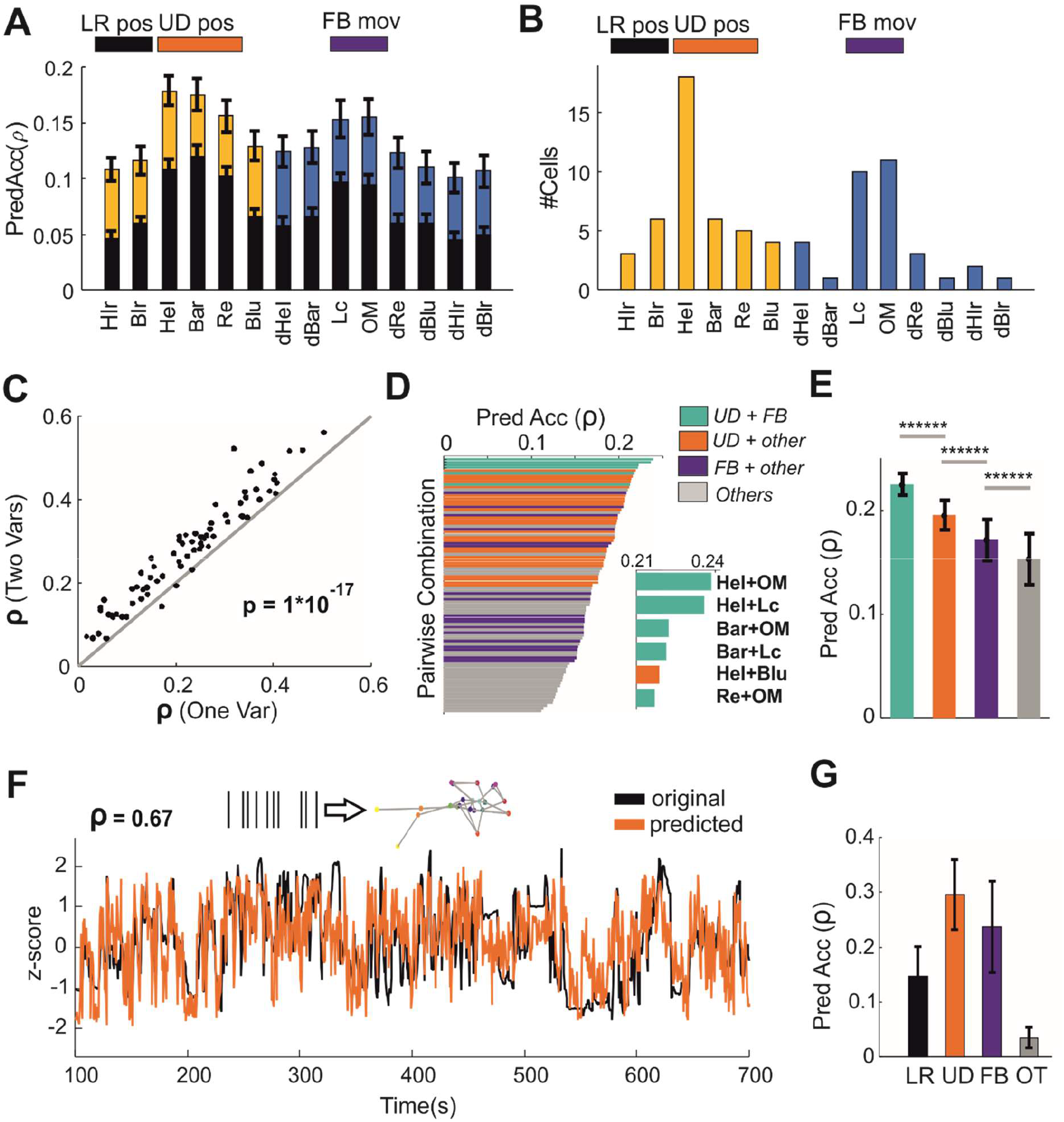
**A-B)** Same as Fig.4A-D but here we report results based on prediction accuracy instead of Mutual Information. **C-E**) Same as Fig.3E-G but for animals recorded under photopic light. **F-G)** Same as Fig.2H-I but for animals recorded under photopic light.

**Supplementary Figure 5:**
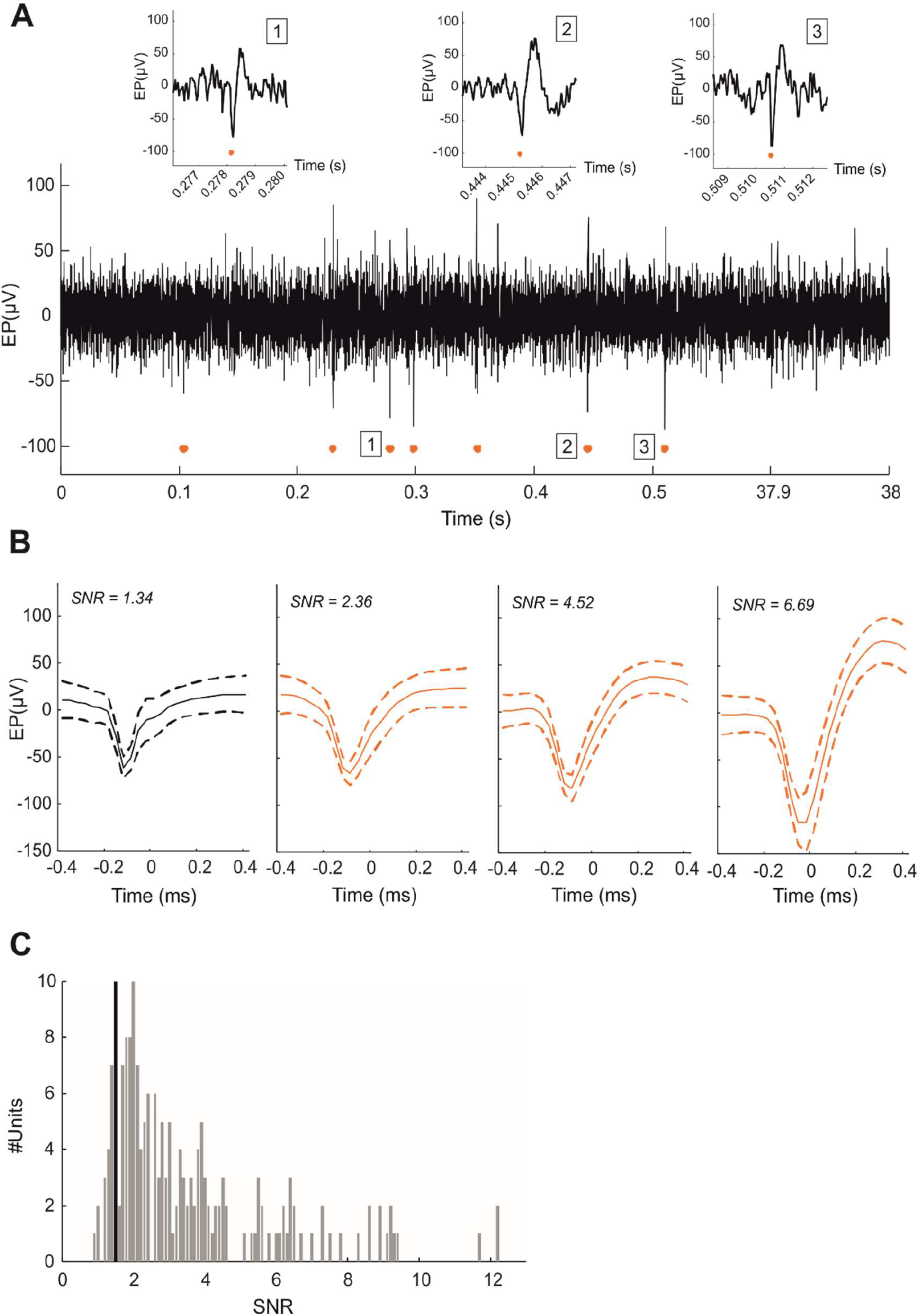
**A)** Representative extracellular recording. Spikes detected are indicated by orange dots and three of them magnified for visual inspection (see insets 1-3). **B)** Representative sample of recorded units. Signal-to-Noise ratio (SNR) is reported at the top of each panel. Only units with SNR>1.5 were used for further analyses (here we excluded the first unit on the left since SNR=1.34). **C)** Distribution of SNR across our dataset (black vertical line indicates the threshold for inclusion).

## Movie Captions

**Movie 1:** Representative movie illustrating 3D reconstruction of behaviour.

**Movie 2:** Illustration of changes in all postures and for locomotion and rearing.

## Notes

### Competing Interest Statement

The authors have declared no competing interest.

### Summary of Updates

I amended the name/affiliations in the text of the manuscript I uploaded a higher resolution pdf (in the previous one the figures were hard to understand due to very low resolution)

## References

1. Land, M., N. Mennie, and J. Rusted, The roles of vision and eye movements in the control of activities of daily living. Perception, 1999. 28(11): p. 1311–28.

2. De Franceschi, G., et al., Vision Guides Selection of Freeze or Flight Defense Strategies in Mice. Curr Biol, 2016. 26(16): p. 2150–4.

3. Hoy, J.L., et al., Vision Drives Accurate Approach Behavior during Prey Capture in Laboratory Mice. Curr Biol, 2016. 26(22): p. 3046–3052.

4. Goodale, M.A. and A.D. Milner, Separate visual pathways for perception and action. Trends Neurosci, 1992. 15(1): p. 20–5.

5. Cao, L. and B. Handel, Walking enhances peripheral visual processing in humans. PLoS Biol, 2019. 17(10): p. e3000511.

6. Miller, E.F., 2nd, A.R. Fregly, and A. Graybiel, Visual horizontal-perception in relation to otolith-function. Am J Psychol, 1968. 81(4): p. 488–96.

7. Ganel, T. and M.A. Goodale, Visual control of action but not perception requires analytical processing of object shape. Nature, 2003. 426(6967): p. 664–7.

8. Jackson, S.R. and M. Husain, Visuomotor functions of the posterior parietal cortex. Neuropsychologia, 2006. 44(13): p. 2589–93.

9. Whitlock, J.R., Navigating actions through the rodent parietal cortex. Front Hum Neurosci, 2014. 8: p. 293.

10. Blot, A., et al., Visual intracortical and transthalamic pathways carry distinct information to cortical areas. Neuron, 2021. 109(12): p. 1996–2008 e6.

11. Wurtz, R.H., et al., Thalamic pathways for active vision. Trends Cogn Sci, 2011. 15(4): p. 177–84.

12. Liang, L., et al., Retinal Inputs to the Thalamus Are Selectively Gated by Arousal. Curr Biol, 2020. 30(20): p. 3923–3934 e9.

13. Schroder, S., et al., Arousal Modulates Retinal Output. Neuron, 2020. 107(3): p. 487–495 e9.

14. Erisken, S., et al., Effects of locomotion extend throughout the mouse early visual system. Curr Biol, 2014. 24(24): p. 2899–907.

15. Ito, S., D.A. Feldheim, and A.M. Litke, Segregation of Visual Response Properties in the Mouse Superior Colliculus and Their Modulation during Locomotion. J Neurosci, 2017. 37(35): p. 8428–8443.

16. Roth, M.M., et al., Thalamic nuclei convey diverse contextual information to layer 1 of visual cortex. Nat Neurosci, 2016. 19(2): p. 299–307.

17. Niell, C.M. and M.P. Stryker, Modulation of visual responses by behavioral state in mouse visual cortex. Neuron, 2010. 65(4): p. 472–9.

18. Keller, G.B., T. Bonhoeffer, and M. Hubener, Sensorimotor mismatch signals in primary visual cortex of the behaving mouse. Neuron, 2012. 74(5): p. 809–15.

19. Saleem, A.B., et al., Integration of visual motion and locomotion in mouse visual cortex. Nat Neurosci, 2013. 16(12): p. 1864–9.

20. Aydin, C., et al., Locomotion modulates specific functional cell types in the mouse visual thalamus. Nat Commun, 2018. 9(1): p. 4882.

21. Nath, T., et al., Using DeepLabCut for 3D markerless pose estimation across species and behaviors. Nat Protoc, 2019. 14(7): p. 2152–2176.

22. Dunn, T.W., et al., Geometric deep learning enables 3D kinematic profiling across species and environments. Nat Methods, 2021. 18(5): p. 564–573.

23. Zhang, L., et al., Animal pose estimation from video data with a hierarchical von Mises-Fisher-Gaussian model. Proceedings of Machine Learning Research, 2021. 130: p. 2800–2808.

24. Storchi, R., et al., A High-Dimensional Quantification of Mouse Defensive Behaviors Reveals Enhanced Diversity and Stimulus Specificity. Curr Biol, 2020. 30(23): p. 4619–4630 e5.

25. Chen, T., Guestrin, C., *XGBoost:* A *Scalable Tree Boosting System,* in *KDD ’16*: Proceedings of the 22nd ACM SIGKDD International Conference on Knowledge Discovery and Data Mining. 2016. p. 785–794.

26. Breiman, L., Random Forests. Machine Learning 2001. 45: p. 5–32.

27. Newman, M.E., Finding community structure in networks using the eigenvectors of matrices. Phys Rev E Stat Nonlin Soft Matter Phys, 2006. 74(3 Pt 2): p. 036104.

28. Stringer, C., et al., Spontaneous behaviors drive multidimensional, brainwide activity. Science, 2019. 364(6437): p. 255.

29. Musall, S., et al., Single-trial neural dynamics are dominated by richly varied movements. Nat Neurosci, 2019. 22(10): p. 1677–1686.

30. Bennett, C., S. Arroyo, and S. Hestrin, Subthreshold mechanisms underlying statedependent modulation of visual responses. Neuron, 2013. 80(2): p. 350–7.

31. Mimica, B., et al., Efficient cortical coding of 3D posture in freely behaving rats. Science, 2018. 362(6414): p. 584–589.

32. Vinck, M., et al., Arousal and locomotion make distinct contributions to cortical activity patterns and visual encoding. Neuron, 2015. 86(3): p. 740–54.

33. Steriade, M., D.A. McCormick, and T.J. Sejnowski, Thalamocortical oscillations in the sleeping and aroused brain. Science, 1993. 262(5134): p. 679–85.

34. McCormick, D.A., D.B. Nestvogel, and B.J. He, Neuromodulation of Brain State and Behavior. Annu Rev Neurosci, 2020. 43: p. 391–415.

35. Liu, D. and Y. Dan, A Motor Theory of Sleep-Wake Control: Arousal-Action Circuit. Annu Rev Neurosci, 2019. 42: p. 27–46.

36. Reimer, J., et al., Pupil fluctuations track fast switching of cortical states during quiet wakefulness. Neuron, 2014. 84(2): p. 355–62.

37. Lee, A.M., et al., Identification of a brainstem circuit regulating visual cortical state in parallel with locomotion. Neuron, 2014. 83(2): p. 455–466.

38. McGinley, M.J., S.V. David, and D.A. McCormick, Cortical Membrane Potential Signature of Optimal States for Sensory Signal Detection. Neuron, 2015. 87(1): p. 179–92.

39. Storchi, R., et al., Modulation of Fast Narrowband Oscillations in the Mouse Retina and dLGN According to Background Light Intensity. Neuron, 2017. 93(2): p. 299–307.

40. Storchi, R., et al., Melanopsin-driven increases in maintained activity enhance thalamic visual response reliability across a simulated dawn. Proc Natl Acad Sci U S A, 2015. 112(42): p. E5734–43.

41. Saleem, A.B., et al., Subcortical Source and Modulation of the Narrowband Gamma Oscillation in Mouse Visual Cortex. Neuron, 2017. 93(2): p. 315–322.

42. Lucas, R.J., et al., Can We See with Melanopsin? Annu Rev Vis Sci, 2020. 6: p. 453–468.

43. Bouvier, G., Y. Senzai, and M. Scanziani, Head Movements Control the Activity of Primary Visual Cortex in a Luminance-Dependent Manner. Neuron, 2020. 108(3): p. 500–511 e5.

44. Dipoppa, M., et al., Vision and Locomotion Shape the Interactions between Neuron Types in Mouse Visual Cortex. Neuron, 2018. 98(3): p. 602–615 e8.

45. Leong, A.T.L., et al., Optogenetic fMRI interrogation of brain-wide central vestibular pathways. Proc Natl Acad Sci U S A, 2019. 116(20): p. 10122–10129.

46. Papaioannou, J.N., Electrical stimulation of vestibular nuclei: effects on light-evoked activity of lateral geniculate nucleus neurones. Pflugers Arch, 1972. 334(2): p. 129–40.

47. Sherman, S.M. and R.W. Guillery, The role of the thalamus in the flow of information to the cortex. Philos Trans R Soc Lond B Biol Sci, 2002. 357(1428): p. 1695–708.

48. Swanson, L.W., O. Sporns, and J.D. Hahn, The network organization of rat intrathalamic macroconnections and a comparison with other forebrain divisions. Proc Natl Acad Sci U S A, 2019. 116(27): p. 13661–13669.

49. Bickford, M.E., et al., Retinal and Tectal “Driver-Like” Inputs Converge in the Shell of the Mouse Dorsal Lateral Geniculate Nucleus. J Neurosci, 2015. 35(29): p. 10523–34.

50. Sokhadze, G., et al., The organization of cholinergic projections in the visual thalamus of the mouse. J Comp Neurol, 2021.

51. Shang, C., et al., Divergent midbrain circuits orchestrate escape and freezing responses to looming stimuli in mice. Nat Commun, 2018. 9(1): p. 1232.

52. Parent, M. and L. Descarries, Acetylcholine innervation of the adult rat thalamus: distribution and ultrastructural features in dorsolateral geniculate, parafascicular, and reticular thalamic nuclei. J Comp Neurol, 2008. 511(5): p. 678–91.

53. Kaplan, H.S. and M. Zimmer, Brain-wide representations of ongoing behavior: a universal principle? Curr Opin Neurobiol, 2020. 64: p. 60–69.

54. Dadarlat, M.C. and M.P. Stryker, Locomotion Enhances Neural Encoding of Visual Stimuli in Mouse V1. J Neurosci, 2017. 37(14): p. 3764–3775.

55. Meyer, A.F., J. O’Keefe, and J. Poort, Two Distinct Types of Eye-Head Coupling in Freely Moving Mice. Curr Biol, 2020. 30(11): p. 2116–2130 e6.

56. Salinas, E. and P. Thier, Gain modulation: a major computational principle of the central nervous system. Neuron, 2000. 27(1): p. 15–21.

57. Eren Sezener, A.G.-B., Dimitar Kostadinov, Maxime Beau, Sanjukta Krishnagopal, David Budden, Marcus Hutter, Joel Veness, Matthew Botvinick, Claudia Clopath, Michael Häusser, Peter E. Latham, A rapid and efficient learning rule for biological neural circuits. bioRxiv, 2021.

58. Storchi, R., et al., Measuring vision using innate behaviours in mice with intact and impaired retina function. Sci Rep, 2019. 9(1): p. 10396.

59. Peirce, J.W., PsychoPy--Psychophysics software in Python. J Neurosci Methods, 2007. 162(1-2): p. 8–13.

60. Mathis, A., et al., DeepLabCut: markerless pose estimation of user-defined body parts with deep learning. Nat Neurosci, 2018. 21(9): p. 1281–1289.

61. Cootes, T.F., Taylor, C. J., Cooper, D. H., Graham, J., Active Shape Models-Their Training and Application. Computer Vision and Image Understanding, 1995. 61(1): p. 38–59.

62. Tipping, M.E., Bishop, C. M., Probabilistic Principal Component Analysis. Journal of the Royal Statistical Society. Series B (Statistical Methodology), 1999. 61(3): p. 611–622.

63. Panzeri, S., et al., Correcting for the sampling bias problem in spike train information measures. J Neurophysiol, 2007. 98(3): p. 1064–72.

64. XGBoost Documentation. Available from: https://xgboost.readthedocs.io/en/stable/.

65. Benjamin, A.S., et al., Modern Machine Learning as a Benchmark for Fitting Neural Responses. Front Comput Neurosci, 2018. 12: p. 56.

